# moRphology - dEep Learning Imaging Cells (RELIC) - to Differentiate Between Normal and Pathological Kidney Exfoliated Cells

**DOI:** 10.1101/2022.04.19.488847

**Authors:** Abbas Habibalahi, Jared M. Campbell, Saabah B. Mahbub, Ayad G. Anwer, Long T. Nguyen, Anthony J Gill, Muh Geot Wong, Angela Chou, Carol A. Pollock, Sonia Saad, Ewa M. Goldys

## Abstract

Chronic kidney disease (CKD) is characterised by progressive loss of kidney function leading to kidney failure. Significant kidney damage can occur before symptoms are detected. Currently, kidney tissue biopsy is the gold standard for evaluation of renal damage and CKD severity. This study explores how to precisely quantify morphology characteristics of kidney cells exfoliated into urine, with a view to establish a future urine-based non-invasive diagnostic for CKD. We report the development of a novel deep learning method, which was able to discover a RELIC (moRphology dEep Learning Imaging Cells) signature that can differentiate between kidney cells exfoliated in human urine of and CKD patients with varying degree of kidney damage and non-CKD controls. Exfoliated proximal tubule cells (PTCs) originating from kidneys were isolated from the urine of patients with different levels of kidney damage using previously published methods. An advanced combination of artificial intelligence techniques, deep learning, swarm intelligence, and discriminative analysis was used to discover a RELIC signature in brightfield microscopy images of exfoliated PTCs. Kidney damage in the study subjects was characterised by assessing kidney tissues obtained through a nephrectomy or kidney biopsy. A special deep learning algorithm was developed and trained to create a predictive tool. The algorithm was then used to analyse data from patients with normal and fibrotic kidneys. Data were then classified according to different groups (healthy or fibrosis) and clustering of the training and validation cells was determined for model validation. We developed a novel deep learning method, to obtain RELIC signatures and identify specific deep morphological features which can be used to differentiate urinary PTC cells shed by people with CKD (confirmed by tissue histology obtained from an invasive kidney biopsy) from those without CKD, with a discriminatory accuracy of 82%. We identified a RELIC signature which can be used on a collection of bright field images of exfoliated urinary PTCs to create a predictive tool and differentiate between normal and pathological kidney cells. This study, for the first time, provides a proof of concept that urinary exfoliated tubule cells in patients with kidney fibrosis and healthy controls differ in appearance (morphology) as observed under a basic brightfield microscope. The results suggest that morphological signatures of exfoliated PTCs have the potential to serve as a non-invasive marker of kidney fibrosis.

## 1. Introduction

Chronic kidney disease (CKD) is characterized by a progressive loss of kidney function leading to irreversible kidney failure requiring dialysis or kidney transplantation (1). Significant kidney damage can occur prior to demonstrable kidney functional decline and accurate or early assessment of kidney damage provides a greater opportunity to preserve kidney function that translates to a better long-term outcome (2). Invasive kidney biopsy is the gold standard for assessing the severity of tubulointerstitial fibrosis and thus the likelihood of progressive CKD (3, 4) but it can not be routinely used to assess CKD disease progression. CKD is currently assessed by serial measurements of estimated glomerular filtration rate (eGFR) and albuminuria (5). However, both of these measures correlate poorly with kidney interstitial fibrosis, the major hallmark of CKD progression (6). Therefore, developing a sensitive noninvasive diagnostic tool for kidney damage would provide clinically important information in addition to eGFR and albuminuria to improve monitoring and management of progressive CKD.

Kidney cells including podocytes, tubular epithelial cells, urothelial cells, and other cell types are continuously shed into urine even in a healthy state (7) and their condition is likely to reflect the changing physiology of the kidneys in CKD, e.g. tubulointerstitial fibrosis (8). To date, there have been few studies using urinary cells as a potential diagnostic tool to assess kidney functional and structural abnormalities (8). This is largely due lack of sensitive diagnostic methods to identify disease-related properties of these cells.

Cellular morphology is altered by biophysical and environmental factors affecting the cells (9) and it has been successfully deployed to probe subcellular mechanisms (10). Therefore, we hypothesised that specific morphological cell signatures representing cell characteristics related to kidney fibrosis in CKD could be determined. Artificial intelligence (AI) is suitable for this task as it is capable of extracting a significant amount of information in the cell images, and we used it in this work to extract image features (11). AI has been successfully used for applications in precision medicine (12) and to extract clinically relevant information in medical images, which is undetectable by expert inspection. For example, deep learning has been applied to improve the diagnostic value of electrocardiography (ECG), electroencephalography (EEG), magnetic resonance imaging (MRI), and magnetoencephalography (MEG), and in all cases it demonstrated exceptional improvement in consistent and accurate decision making (13).

We hypothesize here that exfoliated kidney cells undergo morphological changes due to kidney damage in CKD patients, which may have potential diagnostic value. To test our hypothesis, we used standard brightfield microscopy to capture morphological properties of exfoliated urinary PTCs from patients with CKD and thus defined a RELIC signature of kidney pathology. Accurate prediction using AI models require a large number of parameters that need to be learned. This is manageable when a large amount of training data is available; however, the use of AI and particularly deep learning for risk prediction or diagnosis is currently challenging when the volume of training data is limited. In this work we developed an accurate cellular RELIC signature based on AI and deep learning which can be derived from a limited data volume. Kidney cells extracted from patients with healthy and fibrotic kidneys were imaged by standard brightfield imaging and then used for model building and data validation. RELIC signatures were derived by an advanced combination deep learning (14), swarm intelligence (15), and discriminative analysis (16). To the best of our knowledge, our study is the first that correlates the morphological signatures of exfoliated kidney PTCs with kidney fibrosis status in CKD.

## 2. Materials and Methods

### 2.1. Patients characteristics

To validate our novel method and examine its potential suitability for detecting differences between CKD patients and controls, we used exfoliated kidney cells (PTCs) extracted from patient’s urine. Local ethics approval was obtained for this study (HREC/17/HAWKE/471). Ten consented patients undergoing nephrectomy or clinically indicated kidney biopsy were recruited for this study. A freshly voided urine sample was collected from the patients before the procedure to extract kidney exfoliated PTCs. Biopsy materials and nephrectomy specimens were subjected to histological analysis.

For histological assessment, kidney tissues were stained with haematoxylin, Periodic Acid - Schiff, and Masson’s trichrome stains, and kidney pathology was assessed blindly by 2 independent pathologists to determine interstitial fibrosis and tubular atrophy (IFTA) and scored according to previous studies (17, 18). We had two groups of patients: group 1 with no detectable kidney pathology (controls referred to as “no damage”; n=3) and group 2 with detectable kidney pathology; IFTA in over 25% of kidney tissue (damage; n=7).

### 2.2. Renal proximal tubule cell isolation from urine

Proximal tubule cells (PTCs) were extracted from the urine (100ml) of the above 2 groups using immunomagnetic separation with antibodies against aminopeptidase N (CD13) and sodium/glucose co-transporter 2 (SGLT2) as we have previously described (8). Briefly, urine was spun at 3000/4000 rpm for 20 min (at 4 °C) to pellet the urinary exfoliated cells, which were then washed twice with phosphate buffered saline (PBS). Cells were incubated with mouse anti-human CD13 antibody (Invitrogen @ 5 µg) and anti-SGLT2 antibody (Abcam @ 2.5 µg of) for 1 hr at 4 °C. CELLection Pan anti-mouse Dynabeads (Invitrogen) were then used according to the manufacturer’s instruction to pull PTCs that were positive for both CD13 and SGLT2. Cells were then eluted from the beads and assessed using brightfield microscopy. In total, 49 cells were assessed from the control group and 87 cells were assessed from patients with CKD to validate our deep learning method.

### 2.3. Confocal microscopy

To confirm specific isolation of PTCs, an aliquot of isolated cells was incubated with antibodies against CD13, SGLT2 and angiotensinogen (AGT) at 4°C overnight as we have previously described (8). They were then washed with PBS and stained with the relevant secondary antibodies. Cells were then washed two more times in ice-cold PBS and stained with Hoechst 33342 (Invitrogen, CA, USA) before being loaded onto 35mm coverslip-bottom dishes for confocal imaging (Olympus FV3000, Shinjuku, Tokyo, Japan). Cells attached with debris or poorly focused were excluded during the imaging process. We captured bright field images from the rest of PTCs.

### 2.4. Cellular imaging by brightfield microscopy

Unstained PTCs were imaged by brightfield microscopy using an Olympus iX83™ microscope. Using a 40× (NA 1.15) objective, bright field images were taken by a Photometrics Prime95B™ sCMOS camera (Sensor: GPixel GSense 144 BSI CMOS Gen IV, Teledyne Photometrics, AZ, US) operating below -30°C to reduce sensor-induced noise. The sensor size was 1200×1200 pixels.

### 2.5. Data analysis

Figure 1 shows the data analysis flowchart to discover and validate the cellular deep morphology signature (DMS) in RELIC. After brightfield imaging, cells were segmented, and images were augmented (19) for artificial dataset expansion (**Supplementary material section 1**). Cell images were then provided to deep learning nets (**Supplementary material section 2**) constructed to extract the deeply learned morphology information. Next, the cell images were divided into the training data set (∼75% of data) and the remaining ∼ 25% of cell images formed a validation set. RELIC was discovered using the training data set (∼ 75% of cells) through iterative application of swarm intelligence (**Supplementary material section 3**) and discriminative analysis. To obtain image features for this study, three CNNs (Net1, Net2, and Net3) were employed. Each CNN produced a separate feature set extracted by each net and then all features were pooled together. These three nets have different structures and resolutions, such choice allows to extract comprehensive image information. Net 1 has 153 convolutional layers with its particular filters obtained from ResNet (20). Net 2 encompassed 22 convolutional layers extracted from GoogLe net (21) and Net 3 has 6 convolutional layers acquired from the Krizhevsky net (22). The last layer of each CNN was finetuned by retraining the final filters of each net by using kidney cell images from this study. Using these three nets, we generated ∼7000 deep features for each cell, forming our dataset. In discriminative analysis, the selected feature subset belonging to patient groups is represented in a discriminative space spanned by canonical variables, which are optimal linear combinations of the selected feature. These canonical variables deliver the highest cluster distinction measured by the Fisher Distance (FD) (23, 24). FD computation was then tracked by the next feature selection cycle conducted by discriminative cluster analysis as well as swarm intelligence. FD was the criterion function of the swarm intelligent technique. The feature subset was improved in an iterative process until the achievement of conversion and no appreciable FD change (convergence) was observed. After RELIC cross-validation based on the validation cellular dataset (∼ 25% of data), the RELIC thus obtained was used to train an SVM classifier (**Supplementary material section 4**) for distinguishing cells by the presence or absence of kidney damage.

**Figure 1.**
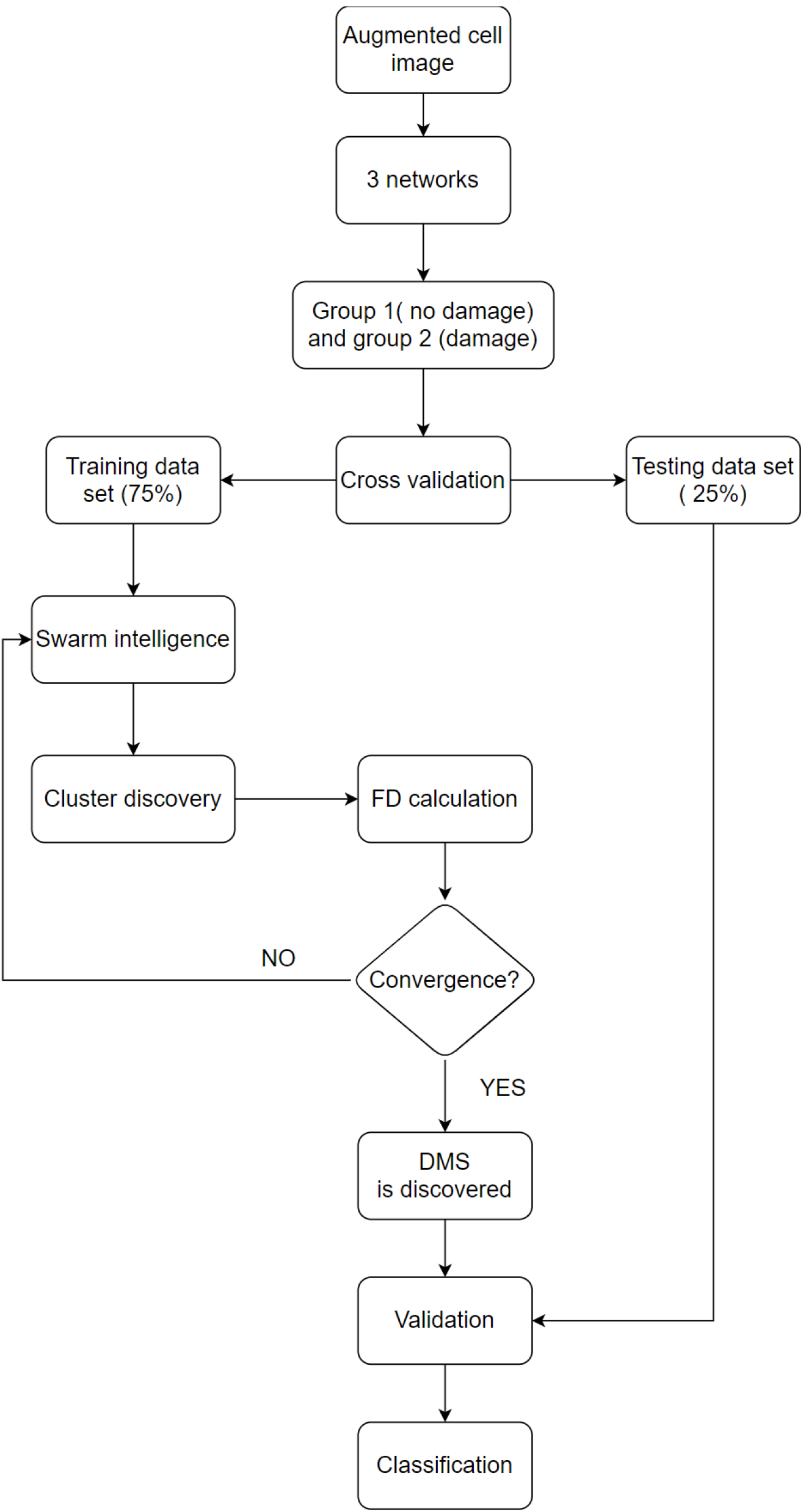
Data analysis methodology flow chart. The procedures and their sequencing are described in Section 2.5.

## 3. Results

### 3.1. Isolated urinary exfoliated cells express proximal tubule cells specific markers

PTCs from the urine were isolated by immune-magnetic separation using SGLT2 and CD13 antibodies as described in the Methods. As shown in (Figure 2), isolated cells express the epithelial marker CD13 as well as kidney proximal tubule-specific markers, SGLT2 and AGT (Figure 2).

**Figure 2.**
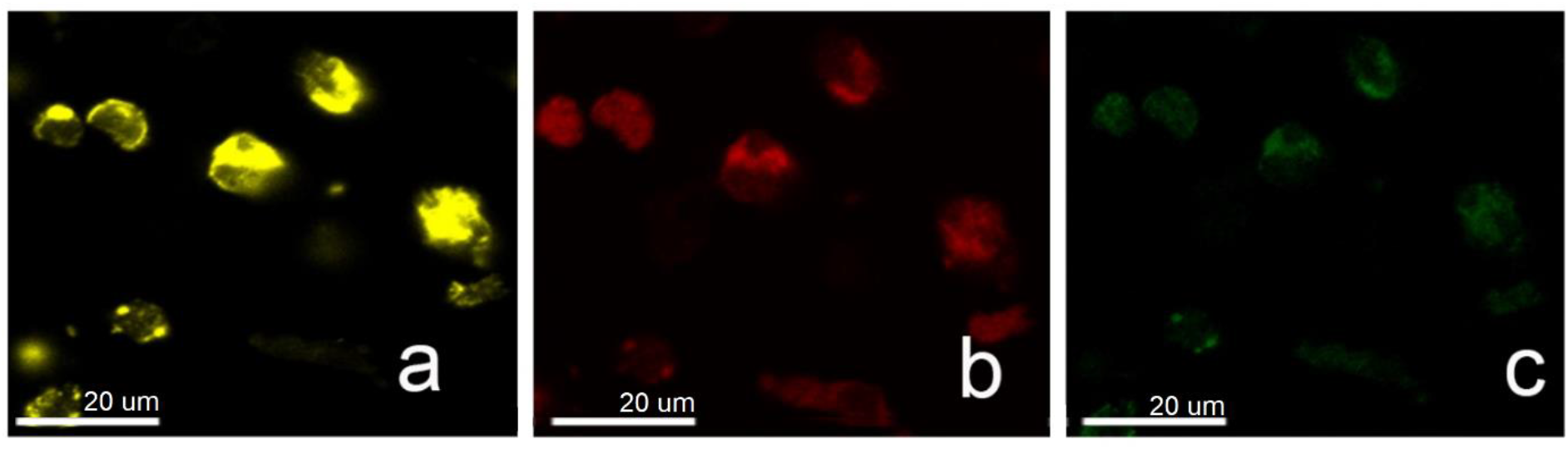
Immunofluorescence staining confirming successful isolation of human urinary PTCs from a patient with CKD. Cells express the epithelial and specific proximal tubule markers (a) CD13, (b) SGLT2 and (c) AGT.

### 3.2. RELIC of exfoliated proximal tubule cells from patients without and with renal pathology

Exfoliated PTCs extracted from individual with healthy kidneys (no damage) and from patients with detectable renal pathology (damage) were analysed. Using the training subset, RELIC was discovered incorporating 7 deeply learned morphological features of PTCs (see section 2.5), and its performance was evaluated. As shown in Figure 3(a), this RELIC could effectively separate cells from patients with and without CKD.

**Figure 3.**
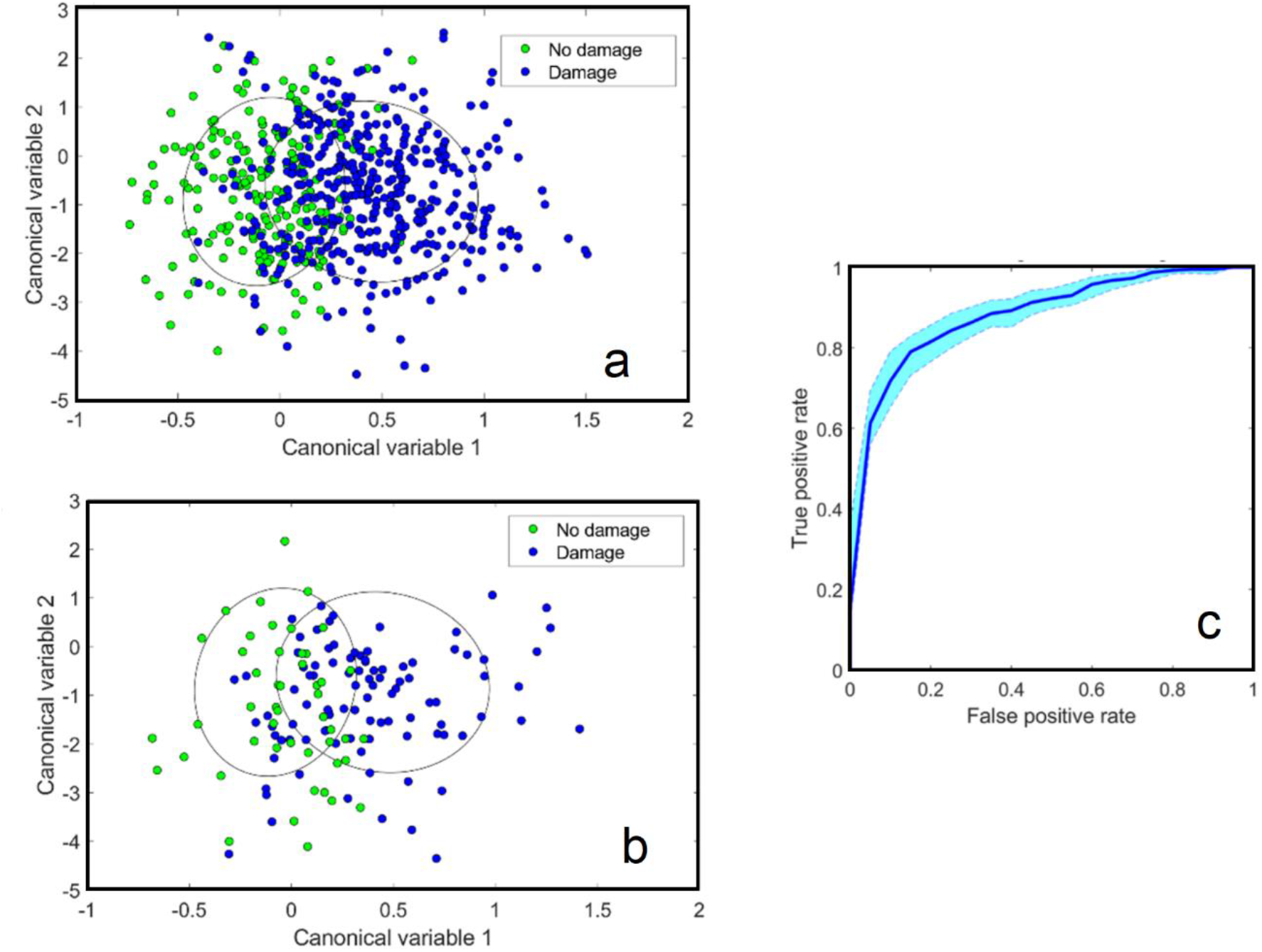
CKD damage classification. (a) Clustering of the training cells from “no damage” and “damage” groups. (b) Clustering of the validation cells from the no damage verse damage groups. (c) ROC curves with 95% confidence interval shown by shaded blue color.

To show the data distribution for each of these disease stage separations, ellipses have been defined for each that encompass one standard deviation from the mean characteristics of cells from healthy and CKD patients. To quantify the overlap of the ellipses, their intersection over union (IoU) values were calculated, ranging from 0% for fully separated to 100% for fully overlapped ellipses, respectively. The IoU values for no damage versus damage patient groups were found to be 18% as demonstrated in Table 1.

**Table 1.** Statistical measures used in this study

| Statistical measure | Value | Statistical measure | Value |
| --- | --- | --- | --- |
| IoU | 18% | True negative rate | 83% |
| Accuracy | 82% | AUC | 0.88±0.02 |
| True positive rate | 79% | P_value (canonical variable 1) | <0.01 |

To evaluate the robustness of the RELIC corresponding to CKD disease, the validation data were employed using cell images, which were put aside during the training process. To this end, the same space created by training data points (shown in Figure 3 (a)) was used; Further, the validation data points were reflected in this space (Figure 3 (b) where ellipses showing the training data distribution from Fig 3 (a) were also drawn. This Figure demonstrates that the clusters of validation data points perfectly coincide with the ellipses created based on the training data points. This proves that our RELIC efficiently separates the validation data points for “damage/no damage” groups in a way similar to the training data points. Additionally, we constructed the SVM classifier (23), which defines cell identity (‘damage/no damage”) based on their RELIC. This is presented in Figure 3 (c) by the receiver operating characteristic (ROC) graphs derived to determine the performance of the classifier. A bootstrapping technique was then used to estimate how accurate are these ROC curves by estimating the sampling error (25). To this end, the cellular data points were randomly resampled 100 times with replacement from our original set of observations, and the corresponding ROC curves were obtained for each classification case, which led to 95% confidence intervals associated with the ROC curve (Table 1). The classification discrimination accuracy, true negative rate and true positive rate were found to be 82% (AUC=0.88±0.02), 83% and 79%, respectively as represented in Table 1.

## 4. Discussion

In this communication, we present a new approach to differentiate between cell groups from a limited number of cell images using deep morphological signatures. We validated our classification method on kidney cells (PTCs) extracted from urine of patients with and without CKD. Our approach is a novel combination of deep learning, swarm intelligence, and discriminative analysis, which can accurately extract and assess morphological properties of cells. First, deep morphological features were extracted using deep learning nets and subsequently swarm intelligence was applied to discover optimized RELIC. Finally, RELIC was employed to train a machine learning classifier (support vector machine) to predict cell labels to damaged kidney cells and healthy cells. Our novel method, RELIC led to an unbiased assessment of cell morphology and was able to accurately identify cells from healthy and damaged kidneys (characterized according to renal pathology) and significantly differentiate between these patients groups. This method requires basic brightfield microscopy imaging only, which will facilitate potential future application of RELIC signatures as a non-invasive diagnostic method for CKD. This richly informative morphological signature distinguished cells derived from patients with kidney pathology from cells from control healthy patients with no pathology with 82% accuracy, 79% true positive rate, and 83% true negative rate (compared with gold standard pathology assessment)). Longitudinal follow-up is required to confirm the clinical applicability of our technique.

As we had 10 samples and in order to minimize the risk of classifier overfitting and to optimize the generalizability, a limited number of deep features were used to train the classifier (7 features), which was less than the root square of augmented cell numbers in each group (26). To evaluate the generalizability our classifier corresponding to kidney fibrosis, the validation was carried out using cell images (20% of data points), which were put aside during the training process (26). In addition, the sampling error was calculated based on standard bootstrapping (25) and the 95% confidence interval of the ROC curve was estimated (AUC confidence interval= 0.02). Bootstrapping proved that the number of observations had a limited effect on our results.

This study shows that we can employ computer vision to discover cell morphological properties characteristic to kidney fibrosis which are not clearly apparent in human visual assessment of microscopic images. RELIC employs deep learning approach capable of extracting specific features which are imperceptible by human visual inspection and understanding such as the specific spatial distribution of image pixel intensities and pixel interrelationships, where there is no current mathematical explanations with which they can be measured (27). This is an advancement over traditional handcrafted features to generate a comprehensive feature bank and independent on operators; however, at the same time, this advantage makes feature interpretability extremely challenging.

We developed a novel deep learning approach to extract morphological cell properties that are specific to kidney fibrosis in CKD by combining advanced artificial intelligent procedures including deep learning, swarm intelligence and discriminative analysis. Our method requires a reduced number of training images compared to traditional deep learning approaches, as it uses pretrained convolutional layers and, employs a limited small number of selected deep features in the final discriminative analysis. Discovering the morphological signature of brightfield cell images using deep features is a novel approach which is presented in this study for the first time. We demonstrated for the first time that deep learning features of urinary exfoliated proximal tubule cells can identify patients with renal pathological changes. Although, a larger patient cohort will need to be assessed prior to the clinical application of this method, the data support the potential of deep learning for non-invasively diagnosing patients with CKD and monitoring its progression. Its ability to diagnose patients at early stages of kidney disease or differentiate patients with different stages of disease needs to be determined.

## Supporting information

supplementary materials in one file

