## supplementary materials in one file for "moRphology - dEep Learning Imaging Cells (RELIC) - to Differentiate Between Normal and Pathological Kidney Exfoliated Cells"

### Supplementary material:

### Supplementary material section 1: Data augmentation

Image augmentation [1] was employed to artificially expand the dataset by adding images equivalent to the original images. To this end, the cell image set was expanded by rotating each cell image by several angles (45$^o$, 90$^o$, 135$^o$, 180$^o$) and reflecting these images with respect to various axes. The image augmentation techniques increase image diversity, increases the number of images and also enhances model performance and generalisability. Convolutional neural networks (Deep convolution neural networks section) trained by image augmentation are able to discover features consistently and irrespective of image location and orientation.

### *Supplementary material section 2:* Deep convolutional neural networks

Deep Convolutional Neural Networks (CNN) is a deep learning strategy for automatic feature identification [2], which is specifically relevant to medical image classification [3]. CNN learns image features through convolutional layers, which are sequentially applied to an input image unlike in conventional methods where features are typically obtained from mathematical descriptions [4]. Convolutional layers consist of several sub-procedures named filters. To extract information, image convolution using collections (arrays) of specific filter is conducted. It is possible to utilise filters published in the literature [5] or they can be developed from the dataset through a learning process [6]. Filter arrays can be trained by assigning random parameters to each filter which are later optimized by comparing predicted class labels and real class labels via the error backpropagated through the network [6].

Filters are convoluted to the input image and moved through the width and height of the input image. The filter output is the new image with reduced size when compared to the input image. Convolutional layers are followed by activation and pooling layers. Activation layers, named "rectified linear unit (ReLU)" replace negative pixels by zeros and keep all positive pixels. Pooling layers cut the spatial size of the input image, which arereplaced by the maximum value in each 3Xe window. Activation and pooling layers make training process more effective by removing negative values, 'downsampling' (making images smaller) and decreasing the number of elements which the network has to learn. Each convolutional layer leads to a modified output image fed as an input to the next convolutional layer. To obtain differential resolution of information corresponding to the image, convolution, ReLU and pooling processes are repeatedly applied to the input image depending on the net structure.

After using several convolutional layers to learn the features, the CNN shifts to classification through a process called "fully connected layers". The use of several convolutional layers and a fully connected classification approach results in a significant number of unknown parameters, which requires a high number of images for optimization [7]. This is extremely challenging for clinical experiments, which typically have a relatively limited numbers of images. Such limited number of available images reduces the accuracy and generalisability, especially with increasing depth (number of layers) [8]. Therefore, it is efficient to use pre-trained convolutional layers already trained for different and unrelated classification tasks, and only fine-tune the last convolutional layer for the specific classification task [8].

### Supplementary material section 3: Swarm intelligence

Swarm intelligence is used in this study to discover the set of best deep morphological features (Number of features = 7). Swarm intelligence simulates the development of naive information-managing cooperating agents in a group [9]. This process aims to obtain a maximized Fisher distance (FD), which is defined as the ratio of between-cluster and within-cluster distances. Swarm intelligence involves multiple agents individually move about in an abstract space particular to the considered problem[10]. As agents move, they endeavour to build up a criterion set depending on the agents location in the developed abstract space. while optimizing the 7 features instantaneously, the space S is all feasible sets of 7 features. The space S considered as a discrete grid created by particular points $N_I = (n_1, n_2, ... n_7)$, where $n_i$ $(i = 1, ... 7)$ is a feature number ranging from 1 to the maximum number of features ( here N= 7000). The grid number of points is $7000^{15}$ and each grid point $N_I$ is characterised by the fisher distance (FD).

Firstly, we arbitrarily chose 100 grid points, $N_{ini}$ from the space S, and these agent locations form the swarm. Next, agents make a move synchronously with their specific route, according to the regulation set on their progress in the space S. This regulation finds the agent and its grid point $N$ while FD is the overall maximum. In addition, the regulation recognizes the maximum FD over this agent's path and the corresponding grid point on this path, $N_{loc}$ . Then, three grid points, $N_{ini}$ , $N_{loc}$ and $N_{max}$ establish what the next movement of the agent is. This spot is the closest grid point to the defined vector $\alpha_1 N_{ini} + \alpha_2 N_{loc} + \alpha_3 N_{max}$ where was defined to be $\alpha_{1,2,3}$ equal 0.15. The agents' movements continue until their trajectories meet at a last grid point, that is the spot of discovered 7 optimized features.

### Supplementary material section 4: Support vector machine classifier

The support vector machine classifier (SVM) was used in this study. An SVM is a robust supervised classification model which is effective for treating sparse data with a limited possibility of overfitting the problem. An SVM generates a hyperplane [11] forming a boundary in a multi-dimensional feature space, and it then categorizes data points into the groups under consideration, depending on which side of the hyperplane they fall into. An SVM identifies the data labels according to a linear predictor function [12]. In this study, the classifier is trained based on the optimal RELIC. The SVM classifier was evaluated based on a cross-validation methodology wherein data points were folded into 10 groups. The classifier was constructed based on 9 folds, and the tenth fold was used for testing and the calculation of accuracy. Further, the receiver operating characteristic (ROC) graph was determined to determine classifier performance [4].
